# A double-blinded randomised placebo-controlled phase II trial to evaluate high dose rifampicin for tuberculous meningitis: a dose finding study

**DOI:** 10.1101/326587

**Authors:** S Dian, V Yunivita, AR Ganiem, T Pramaesya, L Chaidir, K Wahyudi, TH Achmad, A Colbers, L te Brake, R van Crevel, R Ruslami, R Aarnoutse

**Author notes:** Corresponding author. Sofiati Dian MD, Department of Neurology, Faculty of Medicine, Universitas Padjadjaran, Pasteur 38, Bandung, Indonesia (Postal code 40161). /.

## Abstract

**Background:** High doses of rifampicin may help tuberculous meningitis (TBM) patients to survive. Pharmacokinetic-pharmacodynamic evaluations suggested that rifampicin doses higher than 13 mg/kg intravenously or 20 mg/kg orally (as previously studied) are warranted to maximize treatment response.

**Methods:** In a double-blinded, randomised, placebo-controlled phase II trial, we assigned 60 adult TBM patients in Bandung, Indonesia, to standard 450 mg, 900 mg or 1350 mg (10, 20 and 30 mg/kg) oral rifampicin combined with other TB drugs for 30 days. Endpoints included pharmacokinetic measures, adverse events and survival.

**Results:** A double and triple dose of oral rifampicin led to three and five-fold higher geometric mean total exposures in plasma in the critical early days (2±1) of treatment (AUC_0-24h:_ 53·5 mg.h/L vs 170·6 mg.h/L vs. 293·5 mg.h/L, p<0·001), with proportional increases in CSF concentrations and without an increase in the incidence of grade 3/4 adverse events. Six-month mortality was 7/20 (35%, 9/20 (45%) and 3/20 (15%) in the 10, 20 and 30 mg/kg groups, respectively (p=0·12).

**Conclusions:** Tripling the standard dose caused a large increase in rifampicin exposure in plasma and CSF and was safe. Survival benefit with this dose should now be evaluated in a larger phase III clinical trial.

## INTRODUCTION

In 2016, the WHO published data on 10.4 million new tuberculosis (TB) cases and 1·3 million deaths caused by this disease worldwide, making it the leading single infectious disease killer.**(1)** In turn, tuberculous meningitis (TBM) is the most devastating form of TB. It occurs in 1–6% of patients with TB,**(2, 3)** leading to death or neurological disability in more than 30% of affected patients.**(2, 4, 5)**

Antimicrobial treatment for TBM follows the model for pulmonary TB, with intensive and continuation phases of treatment. It adheres to the same first-line TB drugs and dosing guidelines,**(6)** although it is known that some first-line TB drugs, including rifampicin, achieve suboptimal concentrations beyond the blood-brain and blood-cerebrospinal fluid (CSF) barriers. Rifampicin is a crucial TB drug, evidenced by the high mortality rate in TBM patients with resistance to rifampicin. **(7, 8)** As it takes a long time to develop new drugs to treat TB and TBM, it is important to make the best possible use of existing drugs. We performed a series of studies to evaluate higher doses of rifampicin in Indonesian patients with TBM.

A first open-label, randomised phase II clinical trial showed that a 33% higher dose of rifampicin administered intravenously (13 mg/kg iv) for two weeks led to a three-fold higher exposure to rifampicin in plasma and CSF during the first critical days of treatment, and a strong reduction in mortality at six months after the treatment started (adjusted HR 0·42, 95% CI 0·20–0·91).**(9)** We found a clear concentration-effect relationship and derived threshold values for lower mortality, whereas our data suggested that the highest desirable exposures had not been reached yet. **(10)**

However, intravenous rifampicin is not widely available in low to middle-income countries, is expensive, and must be administered by healthcare workers, which is impractical over a longer period of time or unfeasible after discharge from the hospital. Our second open-label pharmacokinetic study was conducted; doses of 17 or 20 mg/kg of oral rifampicin resulted in average total exposures in plasma (area under the concentration versus time curves, AUC_0-24h_) that were approximately similar to the values after 13 mg/kg iv, but average peak plasma concentrations (C_max_) were lower with large interindividual variabilities.**(11)**

Both our first and second clinical trial called for the evaluation of doses of rifampicin higher than 13 mg/kg intravenously and 17–20 mg/kg orally. The current study evaluated the pharmacokinetics, safety/tolerability, and efficacy of higher (up to 30 mg/kg) doses of oral rifampicin as TBM treatment.

## PATIENTS AND METHODS

### Patients

All patients over 14 years old (adults) with clinically suspected meningitis who presented themselves at Hasan Sadikin Hospital, Bandung, Indonesia (the referral hospital for West Java) between December 2014 and November 2016 had an initial screening that included standard CSF tests, blood measurements, and chest radiography.

TBM was classified as ‘definite’ (microbiologically proven) if either CSF microscopy for acid-fast bacilli, Mycobacterium tuberculosis culture, or PCR results were positive. Based on prior evaluation of CSF characteristics of definite and clinically suspected cases in Bandung cohort, patients were classified as having ‘probable TBM’ if they had a CSF/blood glucose ratio <0.5 combined with CSF cells count ≥5cells/μl.**(4)**

Patients with the clinical suspicion of TBM were eligible for the study if they fulfilled the inclusion and exclusion criteria listed in Supplement 1. An HIV test was performed in every patient. Microbiological examinations were performed for cryptococci in HIV-infected patients (India ink microscopy and cryptococcal antigen testing), M. tuberculosis (Ziehl-Neelsen microscopy, culture, and GeneXpert®)**(12)**, and bacterial pathogens (Gram staining). The neurological status of patients was classified according to a modification of the British Medical Research Council (BMRC) grading system as 1 (Glasgow Coma Scale (GCS) 15 with no focal neurological signs), 2 (GCS 11–14 or 15 with focal neurological signs), or 3 (GCS <10).

Written informed consent to participate in the trial was obtained from all patients or from their relatives if the patient could not provide informed consent. If in the latter case a patient regained the capacity to consider participation, he was consulted and informed consent to continue the study was obtained.

### Study design

The overall aim of this study was to find and substantiate the dose of rifampicin to be studied in a larger follow-up trial, based on pharmacokinetic, safety/tolerability and efficacy considerations. The primary objective was to describe the pharmacokinetics of higher doses of rifampicin in TBM patients. The secondary objectives were to evaluate the safety and tolerability and to explore the efficacy of treatment regimens with higher doses of rifampicin.

The study was a double-blinded, randomised, placebo-controlled phase II trial with three parallel arms. Eligible patients were assigned a standard (450 mg, ~10 mg/kg, one active and two placebo tablets), double (900 mg, ~20 mg/kg, two active and one placebo tablets), or triple (1350 mg, ~30 mg/kg, three active tablets) dose of rifampicin for 30 days, in addition to other TB drugs, according to Indonesian national guidelines. Randomisation occurred in variable block sizes and was stratified by BMRC grade. Adjunctive dexamethasone was administered intravenously according to a previous study in Vietnam.**(13)** After 30 dosages, all the patients continued taking the standard TB regimen up to two months, with subsequent daily doses of rifampicin (450 mg) and isoniazid (300 mg) for a minimum of four months. During hospitalisation in the first four weeks of treatment, adherence to the treatment was secured by facility-based Directly Observed Treatment (DOT). After hospitalisation, adherence was monitored through community-based DOT, frequently through close family members, pill counts and the patients’ medication intake diary.

Rifampicin 450 mg (active) tablets and matching placebo tablets were manufactured at PT Kimia Farma, Bandung, Indonesia. The study drugs and other standard TB treatments were administered on an empty stomach. For unconscious patients who could not swallow, the drugs were crushed and delivered through a nasogastric tube.

The study was approved by the Ethical Review Board of the Medical Faculty of Universitas Padjadjaran, Bandung, Indonesia. External monitoring assessed compliance with the study’s protocol and the Data Safety Monitoring Board (DSMB) performed an interim analysis after half of the subjects were included. This trial is registered with ClinicalTrials.gov, number NCT02169882.

### Bioanalysis and pharmacokinetic (PK) assessments

PK sampling was performed twice, at day 2 (±1) during the first critical days of treatment (PK1) and then at day 10 (±1) of treatment (PK2). On each sampling day, serial blood sampling was done just before and at 1, 2, 4, 8 and 12 hours after dosing. CSF samples for PK assessment were collected on both PK sampling days, between 3 and 9 hours after dosing. Patients had an overnight fast before PK sampling and remained fasted until 2 hours after the administration of the study drugs. Blood samples were centrifuged at 3000 rpm for 15 minutes, and plasma and CSF samples were stored at −80°C within 30 minutes of being taken. Analysis of the rifampicin concentrations in plasma and CSF was performed as described previously. [**11**] The PK parameters for rifampicin in plasma were assessed using standard non-compartmental methods in Phoenix WinNonlin v.6.3 (Certara USA Inc., Princeton, NJ) as described previously.**(14)**

### Follow-up

Patients were followed-up on until six months after their treatment started. All patients were hospitalised for a minimum of 10 days, enabling real-time surveillance of any TBM-related and drug-related adverse events in that period. Monitoring of liver transaminases and full blood counts occurred on days 3, 7, 10, 14, 30, 45 and 60. Any further investigations were performed by indication, e.g. a bilirubin test.

Adverse events in these severely ill patients were defined as those possibly/probably related to their treatment (haematological, gastrointestinal and skin disorders). All other adverse events (e.g. new neurological events and respiratory failure) were not incorporated in the assessment of safety and tolerability.

The classification and grading of adverse events was based on the Common Terminology Criteria for Adverse Event (CTCAE version 4.03). Treatment was stopped in patients who had grade 4 adverse events.

Efficacy was assessed by mortality and clinical and neurological responses on days 3, 7, 30, 60 and 180. New neurological events were defined as the occurrence of any of the following: cranial nerve palsy, motor deficits, or seizures. Modified Rankin Scale (MRS) and Glasgow Outcome Scale (GOS) were used to evaluate the clinical and neurological outcomes.

After discharge, survival was monitored by phone calls or home visits for patients who could not be reached via phone. Verbal autopsies were performed via phone calls and no post-mortem autopsy was done.

### Sample size and statistical analysis

Based on a previous clinical trial, the number of participants needed per study arm to estimate the total exposure to rifampicin (AUC_0-24h_) at a dose of 20 mg/kg, using 95% confidence and a margin of error of 20%, was n=18 for days 2±1 and n=16 at day 10±1.**(11)** Considering dropouts, 20 patients were needed for each study arm. This number of participants was also shown to be sufficient to demonstrate a significant difference in AUC_0-24h_ achieved after 10, 20 and 30 mg/kg of rifampicin.

Patient characteristics, rifampicin PK parameters, adverse events were presented descriptively for all patients and by study group. Multiple linear regression analysis was performed to find predictors of PK parameters. The number of patients with plasma AUC_0-24h_ and C_max_ values above the threshold values for lower mortality of 116 h*mg/L and 22 mg/L at days 2±1 (as determined previously)**(10)** were assessed for each group. Data for safety/tolerability and survival were analysed by the intention-to-treat principle. Mortality data were presented as proportions in each group at various subsequent time points, and proportions were compared using the Chi-square test. A Cox regression analysis assessed the effect of rifampicin dose to 6-month mortality in all patients and for culture-confirmed TBM patients only. IBM SPSS Statistics (version 22.0) for Windows was used for the statistical analyses; GraphPad Prism Version 5.03 and statistical software R version 3.4.2 **(15)** were used for obtaining graphs. P values of less than 0.05 were judged significant in all analyses.

## RESULTS

During the study period, out of the 229 patients with suspected TBM, 107 patients were excluded for alternative diagnoses, and the remaining 122 patients were diagnosed with TBM. The 60 TBM patients who fulfilled all eligibility criteria were randomly assigned to each arm (**Figure 1**). Baseline characteristics are depicted in **Table 1**, showing that the patients’ characteristics were equally balanced across the groups, except for fewer males in the 20 mg/kg group (p=0·343) and a higher percentage of HIV positives (p=0·261) and lower CSF protein concentrations (p=0·057) in the 30 mg/kg group. Overall, 53% were male, with a median age of 29·5 years, and 10% of patients had HIV infection.

**Figure1.**
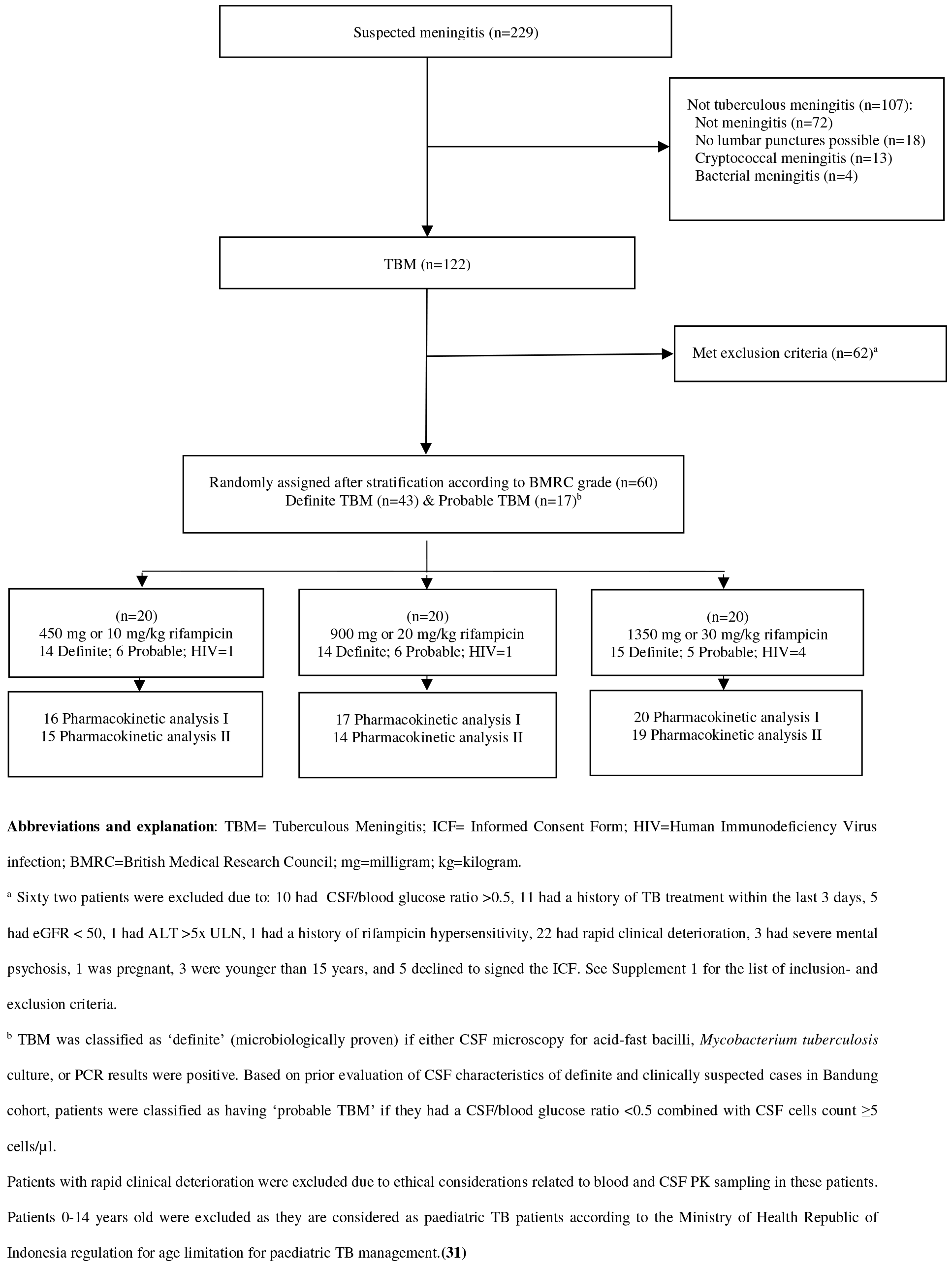
Trial profile.

**Table 1.**
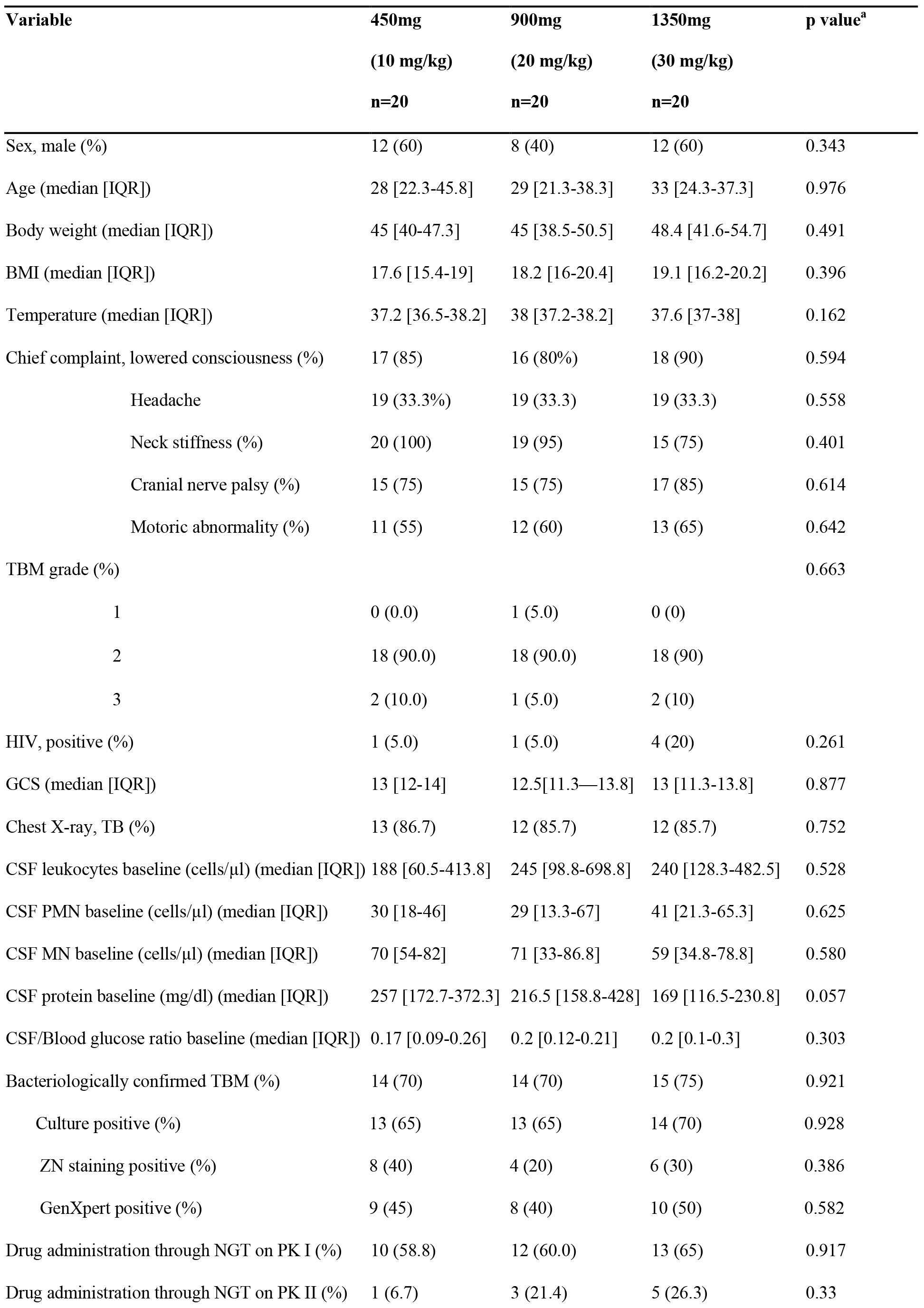
Patients’ characteristics.

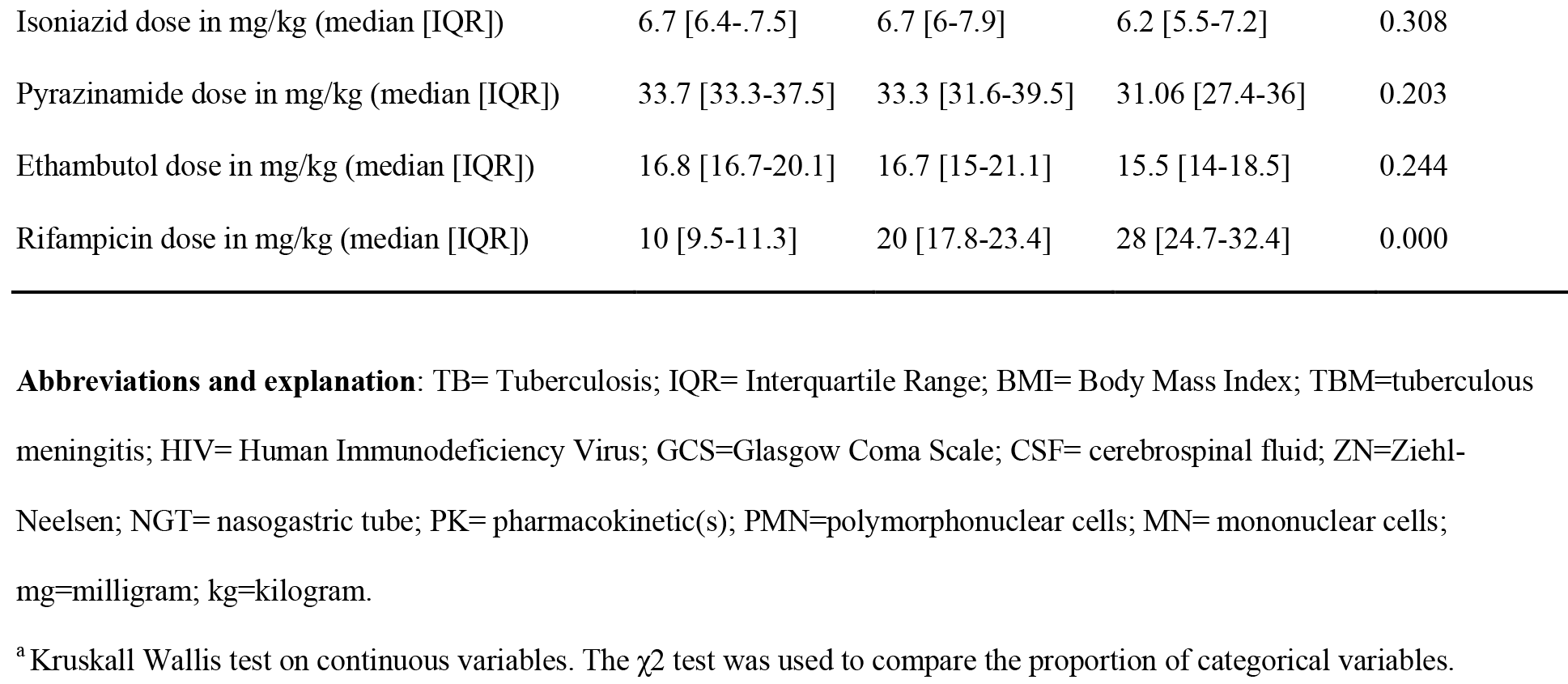

Fifty-three patients had their first PK assessments (PK1) done, whereas 48 patients had a second PK evaluation (PK2). Patients who failed to undergo a PK assessment were those who died or were withdrawn. The study drugs were administered via a nasogastric tube (NGT) for 33 patients (62%) at the first PK assessment and for nine patients (19%) at the second assessment. In some patients, their eventual plasma AUC_0-24h_ and C_max_ values could not be assessed reliably based on sampling at 1, 2, 4, 8 and 12 hours after the dose (**Table 2**) and in some patients concomitant CSF sampling did not take place during the two PK assessments.

**Table 2.** Pharmacokinetic data of rifampicin.

|  | 450 mg<br>(10 mg/kg)<br>GM [range] | 900 mg<br>(20 mg/kg)<br>GM [range] | 1350 mg<br>(30 mg/kg)<br>GM [range] | p value<sup>a</sup> |
| --- | --- | --- | --- | --- |
| **First PK assessment** |  |  |  |  |
| Day (Median [range]) | 1 (1-3) | 1 (1-3) | 1 (1-3) |  |
| AUC<sub>0-24h</sub> (mg.h/L) | 53.5 [13.4-116.5] | 170.6 [26.4-377.1] | 293.5 [190.1-533.3] | 0.000 |
| T<sub>1/2</sub> (h) [median(range)] | 4.9 [2.1-20] | 6.3 [3.7-10.6] | 8.8 [3.7-25.6] | 0.022 |
| C<sub>max</sub> (mg/L) | 7.2 [2.2-14.1] | 18.1 [2-43.6] | 25.5 [11.9-55.5] | 0.000 |
| Vd/F (L) | 59.2 [26.0-217.1] | 47.9 [22.4-516.3] | 58.1 [24.8-146.3] | 0.644 |
| Cl/F (L/h) | 8.4 [3.9-33.6] | 5.3 [2.4-34.1] | 4.6 [2.5-7.1] | 0.020 |
| T<sub>max</sub> (h) (Median [range]) | 3.6 [1-9] | 3.9 [0.9-12] | 3.9 [1-12] | 0.882<sup>b</sup> |
| CSF<sub>3-9h</sub> (mg/L) | 0.28 [0.13-0.48] | 0.59 [0.13-1.7] | 0.74 [0.24-1.20] | 0.000 |
| Time to LP (h:m) (median [range]) | 04:02 [03:02-05:45]<sup>c</sup> | 04:19 [03:02-06:13]<sup>c</sup> | 03:57 [03:15-05:29]<sup>c</sup> |  |
| AUC<sub>0-24h</sub> >116 mg*h/L | 1/11 (9.1%) | 13/15 (86.7) | 17/17 (100%) | 0.000<sup>d</sup> |
| C<sub>max</sub> >22 mg/L | 0/14 | 7/18 (39%) | 14/19 (74%) | 0.000<sup>d</sup> |
| **Second PK assessment** |  |  |  |  |
| Day (Median [range]) | 10 (9-11) | 10 (9-11) | 10 (9-11) |  |
| AUC<sub>0-24h</sub> (mg.h/L) | 39.23 [17.41-66.53] | 144.15 [86.25-241.67] | 187.89 [41.61-392.14] | 0.000 |
| T<sub>1/2</sub> (h) | 2.09 [1.27-7.75] | 3.12 [2.02-6.10] | 3.27 [1.65-7.39] | 0.008 |
| C<sub>max</sub> (mg/L) | 7.55 [4.45-14.3] | 21.89 [13.28-32] | 25.14 [9.9-54.78] | 0.000 |
| Vd/F (L) | 34.57 [16.30-80.66] | 28.21 [17.20-66.77] | 34.47 [19.27-77.37] | 0.464 |
| Cl/F (L/h) | 11.47 [6.72-25.85] | 6.24 [3.72-10.44] | 7.19 [3.44-32.44] | 0.004 |
| T<sub>max</sub> (h) (Median [range]) | 2.72 [1-4] | 2.92 [1-8] | 3.59 [1-12] | 0.768<sup>b</sup> |
| CSF<sub>3-9h</sub> (mg/L). | 0.23 [0.13-0.63] | 0.39 [0.13-0.82] | 0.51 [0.13-1.18] | 0.008 |
| Time to LP (h:m) (median [range]) | 03:41 [03:02-06:00]<sup>c</sup> | 03:51[03:28-05:52]<sup>c</sup> | 04:02 [03:17-05:57]<sup>c</sup> |  |

**Abbreviations and explanation:** AUC<sub>0-24h</sub>= area under the time-concentration up to 24 h after dose; GM= Geometric Mean; C<sub>max</sub>= maximum plasma concentration; T<sub>max</sub>= time to C<sub>max</sub>; CSF= cerebrospinal fluid; CSF<sub>3-9h</sub> = concentration in cerebrospinal fluid between 3h and 9h after drug administration; Vd/F= apparent volume of distribution; Cl/F= apparent total clearance; T<sub>1/2</sub>= Half-life; h= hour; m=minutes; L=litre; mg=milligram; kg=kilogram; LP= lumbal puncture.
Data are number (%) or geometric mean (range).
<sup>a</sup> One-way ANOVA (analysis of variance) on log-transformed PK data, with Tukey's HSD as post-hoc test.
<sup>b</sup> Kruskal Wallis test.
<sup>c</sup> CSF samples had to be taken between 3 and 9 h after the dose and were mostly taken around 4 h after the dose.
<sup>d</sup> Threshold values for reduced mortality based on previous work.(10)
PK1 was at day 2 ±1, PK2 was at day 10±1. AUC<sub>0-24h</sub> values were estimated in 43 patients both in PK1 and PK2. The rest could not be estimated because the elimination rate constant could not be assessed reliably with sampling at 1, 2, 4, 8 and 12h after dosing, preventing extrapolation of exposures beyond the last measurable concentration. C<sub>max</sub> could not be assessed in 7 patients.

A higher dose of oral rifampicin resulted in higher geometric mean AUC_0-24h_ values, i.e. three-fold higher for the 20 mg/kg group and five-fold higher for the 30 mg/kg group, whereas plasma C_max_ and CSF concentrations showed a more proportional increase upon increasing the rifampicin dose, both in PK1 and PK2 **(Table 2)**.

Large interindividual variability was observed in plasma rifampicin AUC_0-24h_ and C_max_ values and in CSF concentrations. For example, the differences between the minimum and maximum plasma AUC_0-24h_ values at the first PK assessment were 9-fold for the 10 mg/kg group, 14-fold for the 20 mg/kg group, and 3-fold for the 30 mg/kg group (**Table 2**, **Figure 2**). In the multivariate analysis, the dose administered was the only predictor of AUC_0-24h_ and C_max_ in plasma and concentrations in CSF, whereas gender, age and BMRC grade were not significant predictors (data not shown). The number of patients with drug administration via an NGT was not significantly different between the three study groups and did not predict AUC_0-24h_, C_max_ or CSF concentrations either. AUC_0-24h_, C_max_ values in plasma and CSF concentrations were all highly and positively correlated to each other. For example, plasma AUC_0-24h_ and CSF concentrations correlated with a correlation coefficient of 0·7 (Spearman’s rho, p= <0·01).

**Figure 2.**
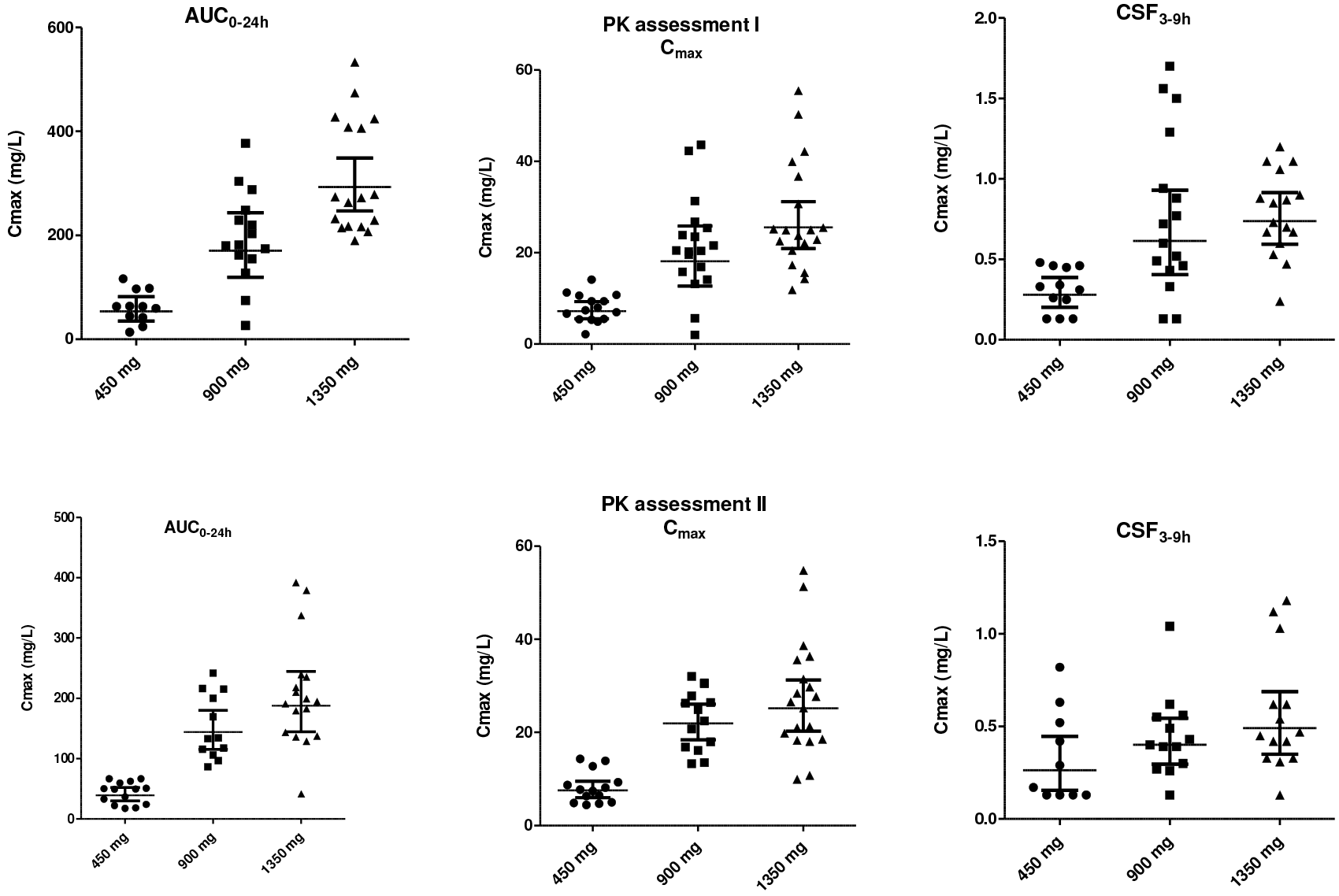
Distribution of rifampicin area under the time-concentration curve up to 24 h after the dose (AUC_0-24h_) and maximum concentration (C_max_) in plasma, and concentration in CSF at 3–9h after the dose, at the first and second pharmacokinetic assessments. Lines represent Geometric Mean (GM) and 95% Confidence Interval (CI).

Comparison of the AUC_0-24h_ values at the first and second PK assessment showed a significant decrease of 33% at the second assessment, p=0·014 (all groups combined), with decreases of 27%, 16%, and 36% in the 10, 20 and 30 mg/kg groups respectively (p=0·21, 0 003, and 0·004, paired T-test on log-transformed AUC_0-24h_ values). Similarly, CSF concentrations decreased from day 2 to day 10 of the treatment, but plasma C_max_ values did not (**Table 2**).

Adverse events during the 30 days after start of study drugs were equally distributed over the groups (**Table 3**). The majority of adverse events was mild. Hepatotoxicity was the most common grade 3 adverse event. All patients with grade 3 adverse events continued treatment and those with grade 3 hepatotoxicity had normal transaminases after a median of 20 days (IQR 7 5–52) of continued treatment without interruption or dose changes of rifampicin. One patient in the 30 mg/kg group had grade 4 hepatotoxicity at day 13 of treatment, as reflected in hyperbilirubinemia of >10 times ULN and transaminases three times the ULN. All TB drugs were interrupted, the dose of rifampicin was unblinded, and the higher rifampicin dose was replaced by a standard dose. The blood total bilirubin had decreased to two times ULN, and transaminases were normalised by day 26 of treatment. Rifampicin AUC_0-24h_ and C_max_ values in those with and without grade 3 or 4 adverse events were not significantly different (p= 0·325 and 0·772).

**Table 3.** Safety and tolerability.

|  |  | 450 mg (10 mg/kg) | 900 mg (20 mg/kg) | 1350 mg (30 mg/kg) |  |
| --- | --- | --- | --- | --- | --- |
|  | All | n=20 | n=20 | n=20 | p |
|  | N (%) |  |  |  |  |
| All AE | 51 (85) | 17 (85) | 16 (80) | 18 (90) | 0.676 |
| Grade I-II AE | 51 (85) | 17 (85) | 16 (80) | 18 (90) | 0.676 |
| Grade III-IV AE | 15 (25) | 3 (15) | 8 (40) | 4 (20) | 0.503 |
| Specific adverse effects |  |  |  |  |  |
| Purpura |  |  |  |  |  |
| Grade I-II | 1 (1.7) | 0 | 0 | 1 (5) | 0.608 |
| Thrombocytopenia |  |  |  |  |  |
| Grade I-II | 8 (13.3) | 2 (10) | 4 (20) | 2 (10) | 0.686 |
| Leukopenia |  |  |  |  |  |
| Grade I-II | 3 (5) | 0 | 1 (5) | 2 (10) | 0.593 |
| Anemia |  |  |  |  |  |
| Grade I-II | 20 (33.3) | 11 (55) | 6 (30) | 3 (15) | 0.107 |
| Grade III | 1 (1.7) | 0 | 1 (5) | 0 |  |
| Hepatotoxicity |  |  |  |  |  |
| Grade I-II | 26 (43) | 9 (45) | 7 (35) | 10 (50) | 0.824 |
| Grade III/IV | 12 (20) | 3 (15) | 5 (25) | 4 (20) |  |
| Nausea |  |  |  |  |  |
| Grade I-II | 27 (45) | 9 (45) | 8 (40) | 10 (50) | 0.444 |
| Vomitus |  |  |  |  |  |
| Grade I-II | 21 (35) | 8 (40) | 6 (30) | 7 (35) | 0.359 |
| Abdominal discomfort |  |  |  |  |  |
| Grade I-II | 15 (25) | 4(20) | 6 (30) | 5 (25) | 0.189 |
| Diarrhea |  |  |  |  |  |
| Grade I-II | 8 (13.3) | 1 (5) | 3(15) | 4 (20) | 0.602 |
| Pruritus |  |  |  |  |  |
| Grade I-II | 27 (43.3) | 9 (45) | 6 (30) | 12 (60) | 0.442 |
| Grade III | 1 (1.7) | 0 | 1 (5) | 0 |  |
| Rash |  |  |  |  |  |
| Grade I-II | 21 (35) | 5 (25) | 5 (25) | 11 (55) | 0.410 |
| Grade III | 1(1.7) | 0 | 1 (5) | 0 |  |

The administration of higher doses of rifampicin increased the percentage of patients who achieved AUC_0-24h_ and C_max_ plasma threshold values for decreased mortality, as derived from our first clinical trial**(9)** (**Table 2**). Two out of the 60 patients were withdrawn from the study on day 2, based on the clinician’s consideration; one after rifampicin resistance was identified by GenXpert®(20 mg/kg group), and the other because of rapid deterioration (30 mg/kg group) as it was considered unethical to perform multiple blood or CSF sampling in this patient. Only an ‘intention to treat’ analysis was deemed useful for this study, as most deaths occurred before the patients could complete the full ‘per protocol’ intervention during one month. This means that the two withdrawn patients were considered to be survivors in our explorative analysis of efficacy.

Six-month mortality was 32% (95% Confidence Intervals (CI): 19.5%–48.1%), and the majority (12/19, 63%) of deaths occurred in the first two weeks after admission. Causes of death were suspected respiratory failure (n=7), septic shock (n=3), brain herniation (n=3), sudden cardiac event (n=2), pulmonary embolism (n=2) and head injury (n=1). Six-month mortality was non-significantly lower in the 30 mg/kg group both among all patients (p=0·12) and among the cases with definite TBM (p=0·07) (**Table 4**). Cox regression analysis showed that the Hazard Ratio (HR) for administration of the triple dose versus the standard dose was 0·44 with 95% CI of 0·1–1·9, p=0·27, in all TBM patients, whereas it was 0·23 (95%CI 0·03–2·05, p=0·19) among patients with definite TBM (both adjusted for HIV status and GCS, see also Supplement 2).

**Table 4.**
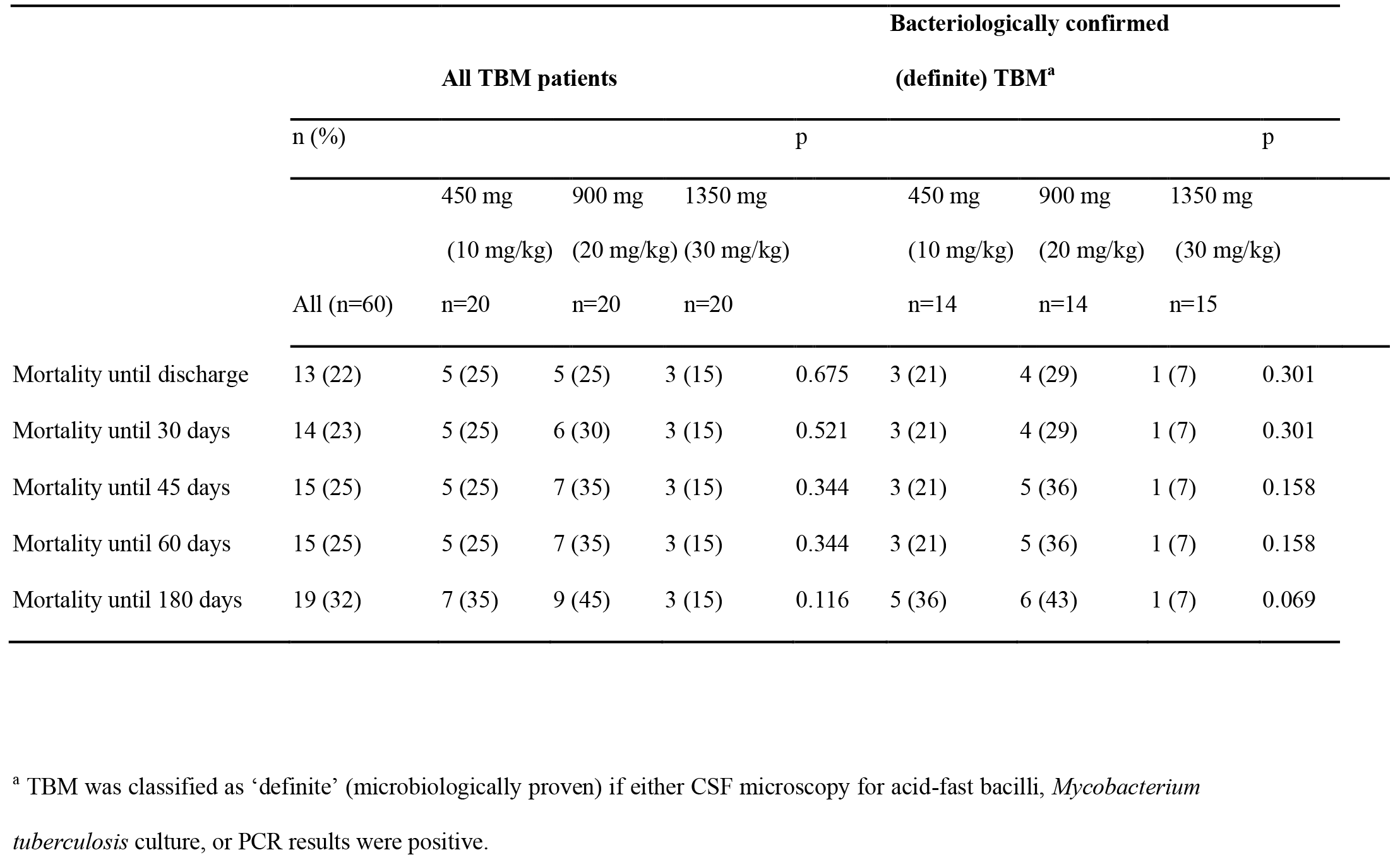
Patients’ cumulative mortality per time point.

New neurological events at day 3, 7, 30, 60 and 180 days, as well as functional outcome measured by MRS and GOS were equally distributed among all three groups of patients (data not shown).

## DISCUSSION

This study revealed that tripling the standard dose of oral rifampicin strongly increased the exposure to this pivotal TB drug in plasma and CSF, did not increase the incidence of grade 3 and 4 adverse events, improved the attainment of previously determined exposure threshold values for lower mortality, and showed a trend for a lower six-month mortality rate among patients with microbiologically proven (definite) TBM.

The standard dose of rifampicin resulted in low average exposures in CSF, around the Minimum Inhibitory Concentration (MIC) of 0·2–0·4 mg/L of this drug for *M. tuberculosis*. **(16, 17** These results are in agreement with data from the literature showing that individual or mean rifampicin CSF concentrations above 1 mg/L are rare.**(17–20)** Due to its protein binding in plasma,**(21)** only 15–20% of rifampicin is available to be transported to other tissues, and penetration to the brain is even more limited.

Fortunately, our intervention with higher doses of rifampicin was effective from a pharmacokinetic point of view. Tripling the rifampicin dose provided a more than proportional increase in AUC_0-24h_ and a proportional increase in plasma C_max_ and CSF concentrations, which were all correlated to each other. A possible explanation for the non-linear pharmacokinetics is the saturation of hepatic extraction or of the excretion of the drug in bile upon increasing the dose.**(22, 23)** The resulting geometric mean AUC_0-24h_ in the 20 mg/kg and 30 mg/kg group in the first critical days of treatment were higher than that of 600 mg i.v. rifampicin in our previous examination**(11)** (170·6 and 293·5 versus 1457h*mg/L); the average C_max_ in the 20 mg/kg group was still lower, but the 30 mg/kg group had a similar C_max_ compared to 600 mg i.v. rifampicin (25·5 versus 24·7mg/L). Our data also showed large inter-individual variability in the rifampicin exposure, which is in agreement with the literature and may be enhanced by pharmacokinetic changes in critically ill patients.**(24, 25** Of note, the lowest observed AUC_0-24h_, C_max_ and CSF concentrations increased with the higher dose (**Table 2**, **Figure 2**); these lowest exposures may be associated with more treatment failures and mortality.

After the first critical days of treatment, we observed decreased exposures to rifampicin after ten days of treatment, which is explained by the auto-inducing enzyme properties of rifampicin. **(22)**

Adverse events after the treatment started were equally distributed among the three rifampicin dose groups. The highest 30 mg/kg daily dose of rifampicin was not associated with an increase in the incidence of severe (grade 3 or 4) adverse events, although it should be noted that the number of patients in this phase II study was relatively small. Hepatotoxicity, as reflected in ALT and AST increases, was the most common grade 3 adverse event and these increases all resolved without any interruption of the study drugs. Thus, based on our data, grade 3 and 4 adverse events and specifically ALT and AST increases did not seem to be dependent on rifampicin dose or exposure. Hepatotoxicity to rifampicin appears to be a hypersensitivity reaction, which is more common with intermittent administration of larger doses. **(26, 27)** Studies of higher doses of rifampicin in pulmonary TB also suggest that hepatotoxicity to rifampicin is idiosyncratic and not dose-related.**(28, 29)**

As to efficacy, tripling the dose of oral rifampicin for a period of 30 days led to an increase in the attainment of plasma AUC_0-24h_ and C_max_ thresholds for decreased mortality and highlighted a trend in survival benefit among those with microbiologically proven (definite) TBM. However, this decrease in mortality was not statistically significant, as our study was not designed and powered to detect a difference in mortality.

The overall aim of this trial was to find and substantiate the dose of rifampicin to be studied in a larger (phase III) follow-up trial. We propose this dose to be 30 mg/kg orally, as this yielded the highest exposure in plasma and CSF, was found tolerable in severely ill TBM patients, and showed a trend for reduced mortality among patients with definite TBM. A study in Africa with a similar 35 mg/kg oral rifampicin dose was associated with an increase in bactericidal activity (**28, 30**) and the same dose achieved a decrease in time to culture conversion in patients with pulmonary TB.**(29)** Ideally, a follow-up trial with rifampicin at a 30 mg/kg dose should be performed amongst TBM patients from various continents.

The strengths of our study are the double-blinding that we applied and the high proportion of bacteriological-confirmed TBM cases (71%). Our study also has limitations. First, the sample size was limited, inherent to the study’s phase II characteristics and objectives, and the study was not powered to compare the efficacy of the three doses of oral rifampicin. As a result, the apparent mortality benefits of high dose oral rifampicin should be interpreted with caution. Second, we could only take single CSF samples for pharmacokinetic purposes, which means that no pharmacokinetic curves could be generated for CSF.

In summary, tripling the dose of oral rifampicin to 30 mg/kg resulted in an increased exposure to rifampicin, both in plasma and CSF, which was not associated with an increase in the incidence of grade 3 and 4 adverse events. The higher 30 mg/kg rifampicin dose resulted in an increase in pharmacokinetic target attainment. In terms of efficacy, further investigation in a larger population is needed to confirm our findings.

## ACKNOWLEDGEMENTS

We thank Ayi Djembarsari and Siti Aminah Soepalarto, Hasan Sadikin Hospital, for accommodating the research; Feby Purnama and Sofia Immaculata, Sri Margi, Shehika Shulda, and Rani Trisnawati for monitoring the patients and data recording; Atu Purnama Dewi for rifampicin bioanalysis and Jessi Annisa for microbiological analysis.

All the authors worked collectively to develop the protocols and methods described in this report. R.R was the principal investigator. ARG and S.D were responsible for the clinical data and the follow-up. Y.V., A.C., and L.t.B performed the pharmacokinetic data analysis under the supervision of R.R. and R.A. S.D. and R.A. performed the statistical analyses with support from K.W. SD performed the literature search and wrote the first draft of the report and all other authors provided contributions and suggestions.

This work was supported by Peer Health (National Academy of Sciences (NAS)-United States Agency for International Development (USAID), USA; the Ministry of Research, Technology and Higher Education, Indonesia (PKSLN grant to T.H.A., R.R., and S.D); the Direktorat Jenderal Pendidikan Tinggi (BPPLN fellowship to S.D.), and Radboud university medical center, The Netherlands (fellowship to S.D).

The authors have no conflicting interests relevant to the study.

Presented in part at the 10^th^ International Workshop on Pharmacology of Tuberculosis Drugs, Atlanta, October 2017.

---

**Supplement 2.**
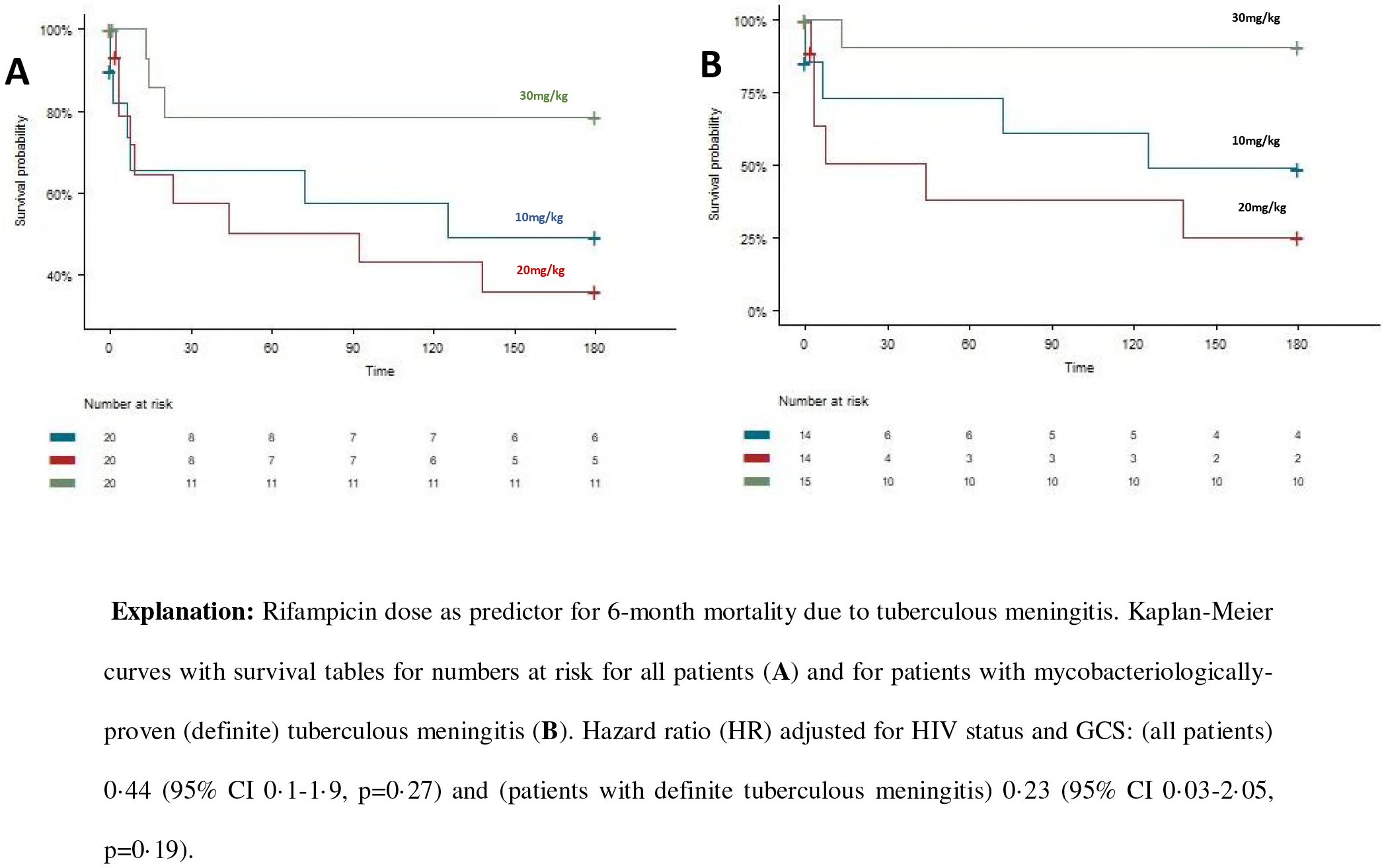
Survival according to rifampicin treatment in all patients and in patients with mycobacteriologically proven (definite) TBM. **Explanation**: Rifampicin dose as predictor for 6-month mortality due to tuberculous meningitis. Kaplan-Meier curves with survival tables for numbers at risk for all patients (**A**) and for patients with mycobacteriologically-proven (definite) tuberculous meningitis (**B**). Hazard ratio (HR) adjusted for HIV status and GCS: (all patients) 0·44 (95% CI 0·1–1·9, p=0·27) and (patients with definite tuberculous meningitis) 0·23 (95% CI 0·03–2·05, p=0·19).

